# Abdominal movements in insect flight reshape the role of non-aerodynamic structures for flight maneuverability I: Model predictive control for flower tracking

**DOI:** 10.1101/2022.06.01.494358

**Authors:** Jorge Bustamante, Mahad Ahmed, Tanvi Deora, Brian Fabien, Thomas L. Daniel

## Abstract

Research on insect flight control has focused primarily on the role of wings. Yet abdominal deflections during flight can potentially influence the dynamics of flight. This paper assesses the role of airframe deformations in flight, and asks to what extent the abdomen contributes to flight maneuverability. To address this, we use a combination of both a Model Predictive Control (MPC)-inspired computational inertial dynamics model, and free flight experiments in the hawkmoth, *Manduca sexta*. We explored both underactuated (*i*.*e*. number of outputs are greater than the number of inputs) and fully actuated (equal number of outputs and inputs) systems. Using metrics such as the non-dimensionalized tracking error and cost of transport to evaluate flight performance of the inertial dynamics model, we show that fully actuated simulations minimized the tracking error and cost of transport. Additionally, we tested the effect of restricted abdomen movement on free flight in live hawkmoths by fixing a carbon fiber rod over the thoracic-abdomen joint. Moths with a restricted abdomen performed worse than sham treatment moths. This study finds that abdominal motions contribute to flight control and maneuverability. Such motions of non-aerodynamic structures, found in all flying taxa, can inform the development of multi-actuated micro air vehicles.

## Introduction

With the exception of feed-forward control, most animal movements are governed by multiple streams of sensory inputs that are processed centrally to coordinate force and torque generation by multiple actuators. Such multi-input, multi-output (MIMO) systems underlie complex motor tasks such as grasping and manipulation as well as myriad modes of terrestrial, aquatic and aerial locomotion (Cowan et al. [2014]). Animal flight is an especially challenging MIMO system that relies on both visual and mechanosensory information processing to coordinate complex wing dynamics for movements ranging from forward flight, to hovering, to tracking moving targets (Cowan et al. [2014], Roth et al. [2016], Taylor and Krapp [2007], Wardill et al. [2017]). The vast majority of literature pertaining to the control and dynamics of animal flight, however, has largely focused on the wing motions for the clear reason that wings generate the lift forces necessary for flight (Dickson et al. [2012], Dickinson [2006]). Yet other body segments may also contribute to movement control and maneuverability by changing the configuration of their position during flight. Inertial redirection of terrestrial locomotion has been observed in lizards as they navigate over rough terrain (Libby et al. [2012]). Likewise the cat righting reflex allows for cats to contour their bodies and land on their feet (Kane and Scher [1969]). For example, tail motions are a key component of the flight control system in birds and bats (Thomas [1996], Su et al. [2012], Gardiner et al. [2011]). In insects, rather than a distinct tail, both the abdomen and legs have been implicated in flight control. For example, leg extension in response to wind gusts in various Hymenoptera have been suggested to serve a role in flight stability (Combes and Dudley [2009]). Similarly, visually driven abdominal flexion and extension is thought to contribute to flight control in a variety of insects including moths (Dyhr et al. [2013]), locusts (Camhi [1970]), and fruit flies (Zanker [1988b,a]). A previous study has shown that while insect flight is inherently pitch unstable, movement of the abdomen yields short-term control in the thoracic pitch of a 2-D simulated butterfly (Jayakumar et al. [2018]). Finally, abdominal undulations (*i*.*e*., periodic abdominal swings coupled to wing inertia) have been demonstrated both experimentally and theoretically to be a mechanism which reduces the overall mean power and mean energy requirements for hovering and forward climbing flight in monarch butterflies (Tejaswi et al. [2021]). These changes in body posture during flight (airframe deformations) may arise either actively (*i*.*e*, movements driven by the insect) or passively (*i*.*e*, respond to external perturbations) and may well contribute to path control.

Given that the abdomen contributes a large proportion of mass for insects (up to 46-67% of body mass for hawkmoths, see Appendix, Table S4), a deeper examination of its role in flight control is warranted. Additionally, understanding the relative roles of wings and abdominal movements in accomplishing specific flight tasks presents an interesting inverse problem: can one predict the forces and torques applied by abdominal motions and wings required to accomplish a specified path? And furthermore, will this system function better in an underactuated state or fully actuated state? Previous literature has addressed the inverse problem of hovering and level forward flight using a combination of genetic algorithm wedded with a simplex optimizer (Hedrick and Daniel [2006]). Moreover, a study that solved the inverse problem by linearizing the dynamical system associated with flight control (Dyhr et al. [2013]) found that flapping flight with active abdominal control operates on the edge of stability.

This paper focuses on flight control in the hawkmoth (*Manduca sexta*) by combining both experimental and computational approaches to address the role of airframe deformations in flight. Computationally, we use an inertial dynamics model of an insect tracking a vertically oscillating signal in which abdominal flexion contributes to the control. Experimentally, we explore this airframe control hypothesis through measurements of free flight behaviors for animals whose abdominal motion is restricted.

We develop an inertial dynamics model inspired by Dyhr et al. [2013] to examine the extent of abdominal contribution in the movement control of a 2D inertial model (an insect) tracking a vertically moving target (a moving flower). We use an approach inspired by Model Predictive Control (MPC) to solve the inverse problem of determining the wing forces and wing and abdominal torques, along with abdominal motions required to achieve a specified flight path (*i*.*e*. to track a vertically moving flower). In doing so, we explore a fully actuated control model (4 controls for 4 degrees of freedom) and an underactuated model (3 controls for 4 degrees of freedom). The degrees of freedom are the two rectilinear directions (*x, y*), and the two rotational directions (*θ, ϕ*). The control freedom is governed by the aerodynamic force (*F*), the direction of the applied aerodynamic force (*α*), the torque applied by the abdomen about the pin joint (*τ*_*abdo*_), and the torque applied by the wings (*τ*_*wing*_). In all cases we analyze both the tracking error of these control strategies as well as a non-dimensional measure of the energy cost associated with each strategy. Interestingly, all MPC solutions were able to solve the flower tracking. However, there are subtle differences in the errors of each approach and substantial differences in the cost of tracking. Additionally, experimentally suppressing abdominal motions in live hawkmoths greatly inhibits flight performance.

### Model formulation and methods

We developed an inertial dynamics model coupled with Monte Carlo simulations to address the question of abdominal contribution to flight maneuverability and flight control. Our model simulates the dynamics of a flying moth as a reduced-order two-mass rigid body system. We used an Euler-Lagrange formulation to generate a system of ordinary differential equations that relate wing forces and body torques to the position and angles of the body in time. Our model includes aerodynamic drag forces on the two body masses (head-thorax, and abdomen mass), and is in the form of a nonlinear, second order system of ordinary differential equations. As we indicate below, we use a method inspired by Model Predictive Control (MPC).

### Insect Geometric Model

The hawkmoth was modeled as two mechanically coupled ellipsoid masses: mass (*m*_1_ and *m*_2_) with associated moments of inertia (*I*_1_ and *I*_2_) (Figure 1b). A pin joint connects the base of the head-thorax mass *m*_1_ to the abdomen mass *m*_2_. The pin joint connection between the two masses was modeled as a damped torsional spring with a spring constant (*κ*) and a torsional damping term (*η*) as shown in Beatus and Cohen [2015]. The externally applied efforts of the system include the aerodynamic force of magnitude (*F*), with a direction (*α*) with respect to the long axis of *m*_1_. Additionally, an abdominal torque at the pin joint (*τ*_*abdo*_), and the wing torque (*τ*_*wing*_) serve as two additional control features.

**Fig. 1.**
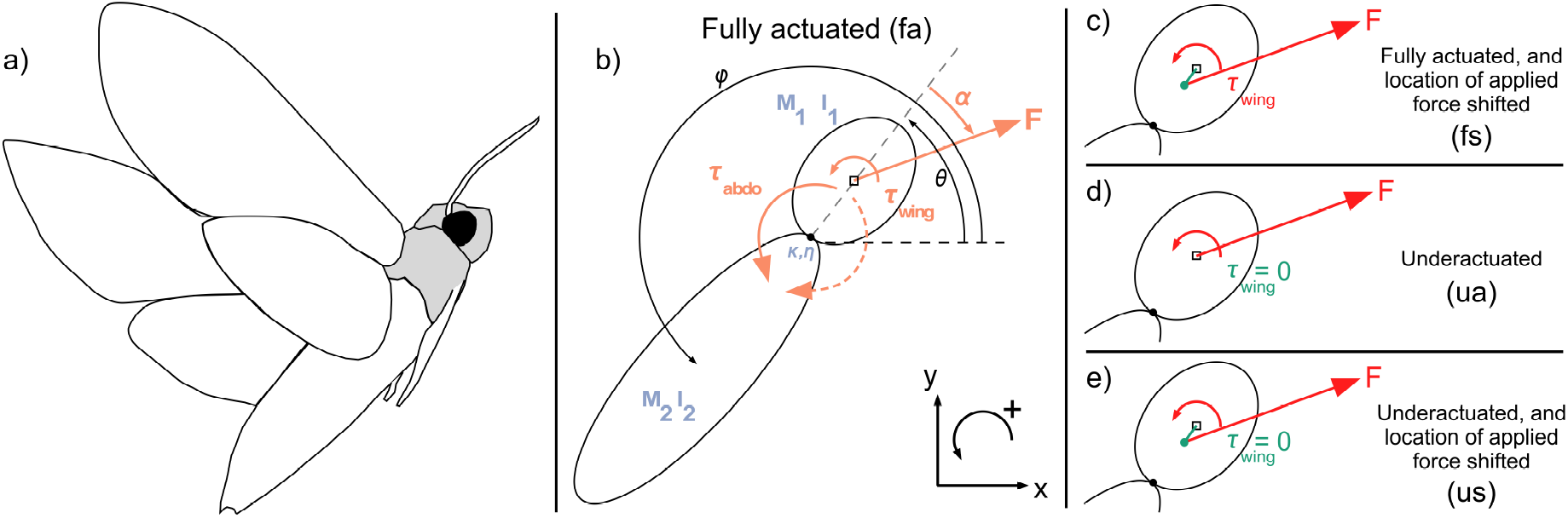
Model basis and modifications. The model has mechanical properties described in blue, and the randomized applied efforts in red. Modifications to the basic model are highlighted in green in (c-e). a) a tracing of a hawkmoth in flight. b) The fully actuated model (“fa”) has two spheroids of prescribed masses and moment of inertia. Reference frame describes positive motion coordinates and rotational motions (counterclockwise). The spheroid of the head-thorax mass is indicated in grey. c) The fully actuated and location of applied force shifted treatment (“fs”) allows for an additional implicit torque on the system. d) The underactuated treatment (“ua”) is identical to the fully actuated treatment (b), with the applied wing torque set to zero. e) The underactuated with location of applied force shifted treatment (“us”) is most similar to treatment (c) but with the applied wing torque set to zero.

The model follows the traditional right-hand coordinate system (*i*.*e*., *x* and *y* are positive going right and upward respectively, Figure 1b). The angular motion associated with the head-thorax mass relative to an inertial coordinate frame is *θ*, while the motion associated with the abdomen mass is *ϕ*, which is also relative to an inertial frame (Figure 1b). To keep consistency with right-handed coordinates, all counter-clockwise rotations are defined as positive. This motion allows for positive pitch to yield a “nose-up” motion.

The model was modified to examine four cases with the modification shorthand in parentheses next to their respective definitions: (1) fully actuated (“fa”, the basic model, Figure 1b), (2) fully actuated with the location of the applied force shifted from the center of mass (“fs”, Figure 1c), (3) underactuated (“ua”, Figure 1d) where the wings do not generate torque, (4) underactuated with the location of the applied force shifted from the center of mass (“us”, Figure 1e). The modifications that include shifting the location of the applied force from the center of the head-thorax mass are included to determine whether the role of implicit torques on the system provide additional stability. By shifting the location of the applied force away from the center of the head-thorax mass, we examine the potential for additional stability using one fewer controller (*i*.*e*. comparing “fs” to “us”). Additionally, these four modified cases were also run with a model where the abdomen mass was decreased by ∼90% to determine the role the applied torques (abdominal and wing torque) contribute to maneuverability. By significantly reducing the mass of the abdomen, we examine to what extent the abdomen plays a role in tracking a vertically oscillating signal, and furthermore determine how flight performance changes.

All model morphometric values and mechanical properties were measured from hawkmoths (*N* = 10, 5 males & 5 females). These specific morphometric values and mechanical properties can be found in the Appendix (Table S2). We used the torsional damping coefficient from Dyhr et al. [2013], and empirically measured the torsional spring constant in this study (see 2.4 below).

### Dynamical Equations

The Euler-Lagrange formulation (Equation 1) yields the equations of motion (see Appendix equations 9-33) necessary to simulate the dynamics of the multi-body model (full derivation in Appendix).

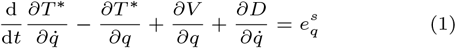

where *T* ^∗^ accounts for the kinetic energies of the system, *V* accounts for the potential energies of the system, *D* accounts for the dissipative energies of the system, and 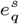 accounts for the work due to external applied efforts. All energies and efforts are with respect to the generalized coordinates. The equations of motion form a coupled system of second-order non-linear ordinary differential equations, which can be separated and solved using an explicit numerical solver we have written in Python (using odeint). All code is available on GitHub.

### Specifying motion and loss function value

We specified a challenging motion for the model to track. We chose prime frequencies as used in Roth et al. [2016], and Sponberg et al. [2015] to determine whether the output response of the model was non-linear. The prescribed goal signal (Equation 2) includes eleven prime number frequencies (see appendix, Table S1), and eleven amplitudes of decreasing magnitude to ensure the velocity of the signal was not increasing as frequency increases.

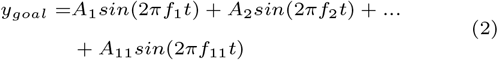

where the amplitudes *A*_*i*_ and frequencies *f*_*i*_ are specified in the Appendix (Table S1) for the *y*_*goal*_. The specified goal motion of the *x*-direction (*x*_*goal*_) is always set to zero. The specified goal angle of the head-thorax mass (*θ*_*goal*_) is always set to *π*/2.

To explore the theoretical control authority, the applied efforts (*F, α, τ*_*abdo*_, *τ*_*wing*_) were randomized using Monte-Carlo methods. The ranges of each applied efforts were defined as follows. The aerodynamic force (*F*) was between 0 and 0.443*kg* · *m* · *s*^−2^ the maximum value is the force required for a hawkmoth to hover in place. The direction of the aerodynamic force (*α*) was between 0 and 2*π* radians to allow for all possible directions of aerodynamic force. The ranges for both the abdominal torque (*τ*_*abdo*_) and the wing torque (*τ*_*wing*_) were between 0 and 0.01*kg* · *m*^2^ · *s*^−2^.

Each individual randomized set of applied efforts yields an individual realization. Each realization was allowed to run for a time period of 20 ms. The time period of 20 ms is half the time of a normal full wing stroke for a hawkmoth. This allowed the model to re-randomize the variables for both the upstroke and the down-stroke, and to further reduce error accumulated over time. 2500 realizations were generated from the randomized set of applied efforts for a given 20 ms time period with the same initial conditions for each particular time period. To simulate a closed-loop system, a loss function (Equation 3) selects one realization out of the 2500 realizations with the lowest loss function value for the particular time period (Figure 2, bottom right blue box).

**Fig. 2.**
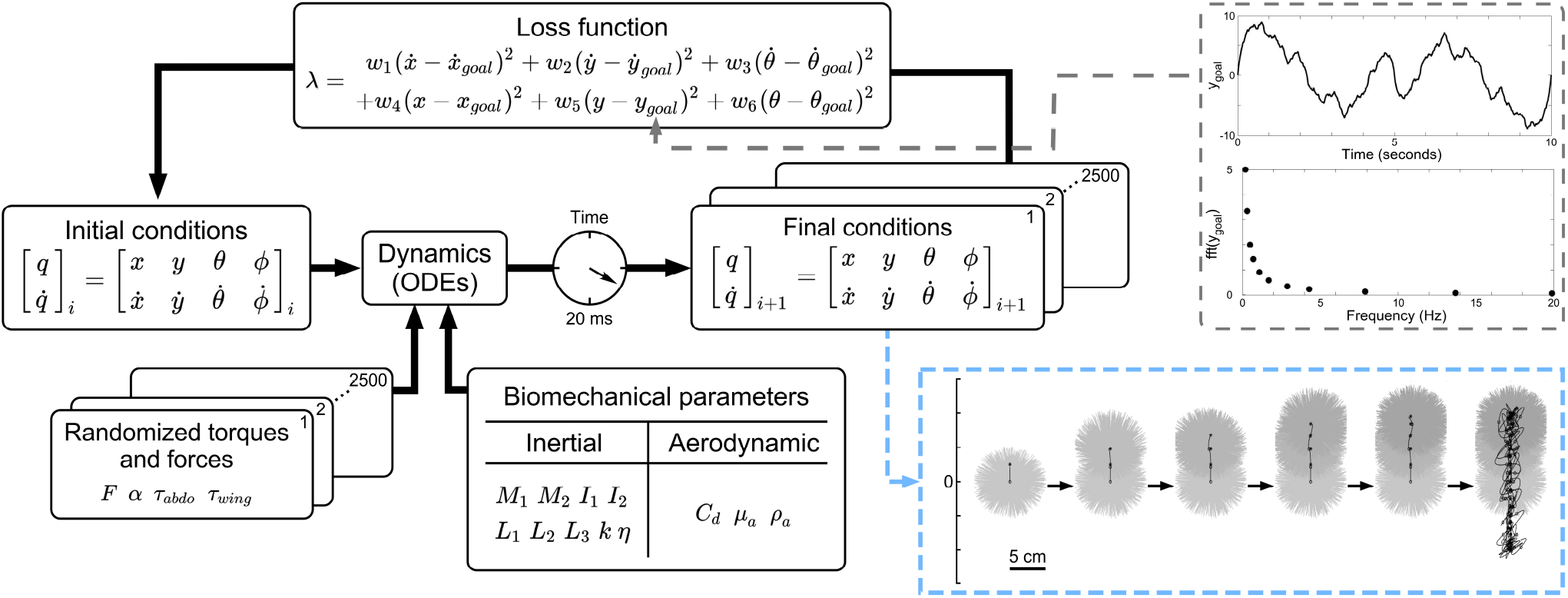
Methods for simulating trajectories. The model has a prescribed set of biomechanical properties: both inertial parameters (masses: *M*_1_, *M*_2_, moments of inertia: *I*_1_, *I*_2_, vector lengths: *L*_1_, *L*_2_, *L*_3_, torsional spring constant: *κ*, and torsional damping coefficient: *η*) and aerodynamic parameters (coefficient of drag: *C*_*d*_, dynamic viscosity of air: *µ*_*a*_, and density of air: *ρ*_*a*_). Each simulation begins with a prescribed set of initial conditions (the set of positions: *q* and their respective velocities: 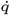). Monte-Carlo methods randomized the applied forces (magnitude of force: *F*, direction of force: *α*) and torques (abdominal torque: *τ*_*abdo*_, and wing torque: *τ*_*wing*_). There are a set of 2500 randomized forces and torques for each time 20 ms time period, yielding the final conditions. The initial conditions, randomized torques and forces, and biomechanical parameters are all passed into the ordinary differential equations (ODEs) of the system (to see the ODEs and their full derivations, see Appendix Equations 30 - 33). A loss function (*λ*) selects the trajectory with the lowest loss function value. The loss function contains various weights (*w*_1−6_, see Appendix Table S3) which penalize deviations from *y* and *x* more than *θ*. The composition of the vertically oscillating signal *y*_*goal*_ is noted in the grey dashed box on the upper right. The time series of the *y*_*goal*_ is displayed for a 10 second time period, the frequency components of this signal are noted in the figure below. The values of amplitude and frequency are noted in Appendix Table S1. The trajectory selected by the loss function uses the values 25% through the trajectory as the new initial conditions for the next 20 ms time period. A visualization of the trajectories and the selection of one trajectory through time is included in the light blue dashed box on the lower right.

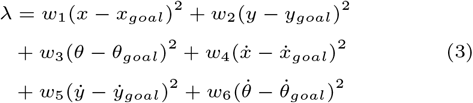

The loss function takes the difference between the prescribed goal value at the end of the 20 ms simulation, and where the realization actually ended up. This is calculated for the head-thorax motion (*θ*), rectilinear position (*x* and *y*), and their respective derivatives(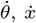, and 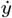). Each term has a prescribed weighting coefficient (*w*_1−6_ see Appendix Table S3) intended to minimize deviations of the head-thorax motion and rectilinear motion.

Drawing inspiration from MPC, we incorporated a receding horizon to minimize the time-accumulated error throughout the 20 ms simulation time. After the loss function selects the realization with the lowest loss function value, the model travels 25% of the selected path. The state variables at this 25% point become the new initial conditions, and we simulate a new batch of 2500 realizations (Figure 2). This method is repeated until the simulation time of 10 seconds is completed.

### Evaluating simulated flight performance

The model simulation flight performance was evaluated using two key metrics: (1) non-dimensionalized tracking error, (2) cost of transport.

We defined the non-dimensionalized tracking error (equation 4) as the rectilinear distance between the the final location of the 20 ms simulation (*x*_sim_, and *y*_sim_) and the goal position. This distance is normalized by dividing the body by length of the organism.

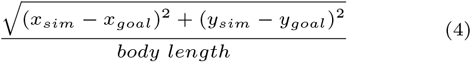

The generalized form of mechanical work is defined in equation 5 as a summation of rectilinear work (work done by the wing force in translational motions), rotational work (work done by the torque resulting from wing forces), and the work due to applied torques (wing torque and abdominal flexion torque).

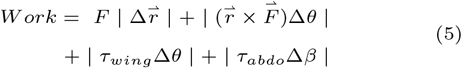

where the first term concerns rectilinear work, and the subsequent three terms describe different rotational work components. *F* is the magnitude of the applied force, 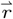 is the generalized positional vector (see appendix), 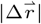 is the distance the simulated moth traveled in each time step of the 20 ms simulation. And the abdominal flexion is defined as *β* and is defined as: *ϕ*_*o*_ - *θ*_*o*_ - *π*. The absolute value of each term in equation 5 represents the energy expenditure of movement in space. Note each model treatment as denoted in Figure 1 may not contain every term in equation 5 for the instantaneous work.

The cost of transport describes the normalized energy expenditure of movement in space. The cost of transport normalizes the mechanical work, as referred to in equation 5, by dividing the product of the weight and distance traveled by the moth at any 20 ms interval as seen in equation 6.

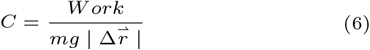

where *m* is the mass of the head-thorax section, and *g* is the acceleration due to gravity.

### Experimental materials and methods

Hawkmoth preparation and abdominal restriction methods

Tobacco hawkmoths (*Manduca sexta*) were collected from a colony at the University of Washington. The moths were maintained on a 12:12 hour light-dark cycle. For the free flight experiment, we used 2-5 days post-eclosion hawkmoths, and specifically selected adults who showed eagerness to feed (actively flew or hovered in front of a red-light headlamp) ∼2 hours after the beginning of the dark cycle.

For the experimental treatment (3 moths, a total of 12 trials), a single carbon fiber rod (0.048-0.095 g) was glued on the dorsal side across the thorax and abdomen joint using superglue to restrict the motion of this joint. Carbon fiber rods for the experimental treatment composed 0.40% +/-0.14% (mean +/-standard deviation) of moth body mass.

A sham treatment was also incorporated in a separate set of moths (4 moths, total of 14 trials). These sham treatment moths had two separate carbon fiber rods glued to the thorax and abdomen respectively (total weight range 0.047-0.067 g). Carbon fiber rods for the sham treatment composed 0.24% +/-0.03% (mean +/-standard deviation) of moth body mass. The purpose of the sham treatment was to account for the weight of the carbon fiber rod. For moths that flew, their last exposure to light was 1.03-4.78 hours (or 62-287 minutes) before flight experiments.

All free flight experiments were performed during the active, night period of hawkmoths including dusk and dawn at 20-25°C. All moths were flower-naíve and had never fed as adults prior to experimentation.

A separate set of 10 hawkmoths (5 males, 5 females) were used to measure the percentage of abdominal mass (as reported in the Introduction). For these measurements, we used 1-3 days post-eclosion hawkmoths. The total mass of the moth was weighed prior to cold anesthetization. After at least 45 minutes of cold anesthetization, the head, thorax, and abdomen were all excised and weighed separately. The value reported in the introduction is the range of percentages of abdominal mass for each individual moth (see Appendix, Table S4).

### Free flight behavioral setup

To document flight behaviors we used an experimental system established by Deora et al. [2021] (see Deora et. al, “Behavioral setup”). Moths flew in an artificial arena (36” x 27” x 36”), darkened, and enclosed from all external lighting. We used an overhead video camera (Basler piA640-210gm GigE) recording at 100 frames per second, 200 *µ*sec exposure under infrared illumination. The enclosure was illuminated with diffuse white light at 0.1 lux with a custom-made LED lightbox (SpyeGrey film, Spyeglass™).

The moths were tasked with feeding from an artificial, 3D printed, funnel shaped, stationary flower (Deora et al. [2021]). We used a 7-component scent mixture which mimicked the scent of hawk moth-pollinated flowers to motivate flight (Campos et al. [2015], Riffell et al. [2013]: 0.6% benzaldehyde, 17.6% benzyl alcohol, 1.8% linalool, 24% methyl salicylate, 3% nerol, 9% geraniol, 0.6% methyl benzoate in mineral oil). A few drops of this scent was placed on filter paper. The filter paper was placed directly above the artificial flower on the ceiling of the artificial enclosure, and covered with a dark cloth so as not to cause a visual distraction for moths during the experiment.

### Evaluation of moth free flight performance

We tracked the moth head using DeepLabCut (DLC, Mathis et al. [2018], see also methods of Deora et al. [2021]) to capture and analyze flight paths of the abdominally restricted and sham treated moths. The DLC model was trained on a previous training data set in Deora et al. [2021] which used similar overhead view of moth feeding from artificial flower. We wrote custom codes (available on GitHub) to analyze these flight tracks. We used several metrics to capture flight dynamics and flight path characteristics like mean velocity, mean acceleration, flight path tortuosity, box dimension as well as analyzed the spectral characteristics of the path. We used a Wilcoxon rank sum test to analyze the differences between abdominally restricted and sham moths across all metrics.

We computed the path tortuosity (equation 7), a metric of how much a moth deviates from a mean or straight flight path to compare flight paths across our treatments. Because moths flight path often cross themselves in this behavioral paradigm, we analyzed tortuosity along smaller segments of the path, essentially creating a sliding window estimate of tortuosity of 40 frames (100 ms, equivalent to 10 wing beats which we felt was an adequate time scale to capture relevant flight trajectory features) for each trajectory and computed the total distance and displacement for each window. We used the following equation to compute tortuosity of each segment (Equation 7), and average across these to compute the mean tortuosity for each flight path. Similarly, we also used fast Fourier transform on each trajectory segment to compute the mean power at various temporal frequencies.

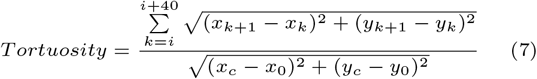

where *i* = [*i, N* − 40] and *N* is the number of frames for any trial. This represents a sliding window of tortuosity estimates. We also used fractal dimensions (box counting dimension) to quantify the complexity (*i*.*e*. jaggedness) of each flight trajectory. Fractal dimensions have been previously used to quantify the geometry of animal paths (Benhamou [2004]). They measure how shapes and patterns scale in comparison to the space they occupy. Hence smaller a fractal value (∼1) means a measured dimension is more similar to a line; higher values (>1.5) means a measured dimension more similar to that of an area. Prior to analysis, each trajectory was saved as an image, so that the space between discrete centroid positions was linearly interpolated. This allowed us to analyze the entire shape of the trajectory irrespective of the moth’s velocity.

### Double-blind scoring of rolling maneuvers

With a single camera system, body pitch and roll are difficult to quantify, but human observers are able to assess the presence or absence of such motions. To determine whether or not there was an increased incidence of rolling in experimental treatment moths relative to sham moths, we used a double-blind study of the moth video trials. Eight volunteers were tasked with the simple binary scoring (yes/no) of whether or not they witnessed rolling in the 26 flight bouts (12 videos of experimental treatment flight bouts, and 14 videos of sham treatment flight bouts). The order of the videos was randomized, and the treatment of the moth was hidden to the volunteer. The metric of evaluation for this scoring was if the incidence of rolls in experimental treatment moths were higher than the incidence of rolls in the sham treatment moths.

### Torsional spring constant measurements

The torsional spring constant was empirically measured for our multibody dynamics model. All torsional spring constant experiments were performed on 1-2 days post-eclosion dead hawkmoths (*N* = 20 moths). There were two groups of dead moths: moths in the freezer (-20°C) for 1 hour before the head was excised (*N* = 10 moths; 5 male, 5 female) and 10 minutes before (*N* = 10 moths; 5 male, 5 female) the head was excised. At the time of head ablation, the wings and legs were also ablated. The scales were removed from the thorax so as to mount the moth into its measurement rig more easily.

Moths were mounted on a micro manipulator with the thorax firmly pierced by two nails to maintain its position during measurements. The abdomen was aligned with a the true vertical as best as possible, and lassoed to the lever arm of a Dual Mode Muscle Lever System (Aurora Scientific, Model 360) firmly with thread.

The Dual Mode Muscle Lever System had the dual task of applying the sinusoidal oscillations and recording the torque that the abdomen exerted back on the lever arm as a response. Calibration of the Dual Mode Muscle Lever System to the corresponding angles and torques was done by tying a rubberband (Sparco Premium Quality Rubber Bands, 1614LB, Size 16, 2 1/2” x 1/16”) to a vertical rod. All data acquisition was performed with a National Instruments DAQ (NI USB-6229) at a sampling frequency of 1000 Hz.

The prescribed sinusoidal signal was 15 seconds long, and composed of one of five driving frequencies (0.2, 1, 5, 10, 20 Hz), with one of three input voltages (2, 5, 9 Volts) which correspond to the different angular sweeps of the lever arm (5.42, 13.56, 24.41 degrees respectively). Each of these 15 signals had three replicates. The order of these 45 trials were randomly permuted for each of the 20 moths to ensure there was no statistical bias or unusual response by the moth abdomen to different driving frequencies and amplitudes.

We recorded the time series for both the abdominal angle and applied torque for these 45 trials. Both time series signals were Fourier transformed. These two Fourier transformed signals each have a real component and an imaginary component. The real components of the two Fourier transformed signals is used to calculate the torsional spring constant:

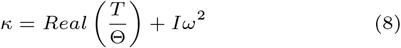

where *T* is the Fourier transform of the torque signal, Θ is the Fourier transform of the angle signal, *I* is the moment of inertia of the moth abdomen, and *ω* is the driving frequency. For the full derivation, see the Appendix.

All instantaneous torsional spring constants (for all trials and moths) were plotted with respect to driving frequency and fit with a second degree polynomial function. The *x*-intercept of this polynomial fit was the torsional spring constant used in all of our simulations. The torsional spring constant value we used in the simulations is 0.0023 *kg* · *m*^2^ · *rad*^−1^ · *s*^−2^ (see Results).

## Results

### Torsional spring constant measurements

Moths which were in the freezer for 10 minutes prior to head excision had a torsional spring constant of 0.0023 +/-0.00072 *kg* · *m*^2^ · *rad*^−1^ · *s*^−2^ (mean +/-standard deviation). Moths which were in the freezer for 1 hour prior to head excision had a torsional spring constant of 0.0034 +/-0.0020 *kg* · *m*^2^ · *rad*^−1^ · *s*^−2^ (mean +/-standard deviation). We decided to use a value within the range of the moths which were in the freezer 10 minutes prior to head excision because such moths have material properties more directly relevant to the live moths.

### Simulated flight: non-dimensional tracking error

Non-dimensional tracking error denotes the 2-dimensional rectilinear distance between the final state of the model in a 20 ms simulation run time, and the goal position normalized to body length (*L*_1_ + *L*_2_, constant for all simulations) of the simulated moth. A Kruskal-Wallis one-way analysis of variance test showed at least one of the groups were significantly different from the other (Kruskal-Wallis, *χ*^2^ = 294.14, *df* = 7, *P* = 2.2e-16, effect size, *η*^2^ = 0.941): the Dunn’s test for multiple comparisons revealed the treatments which had the lowest non-dimensionalized tracking error were the two fully actuated treatments for the regular-sized abdomen, and one underactuated treatment for the reduced mass abdomen (regular-sized abdomen “fa” and “fs” in Figure 3a, and reduced mass abdomen “us”).

**Fig. 3.**
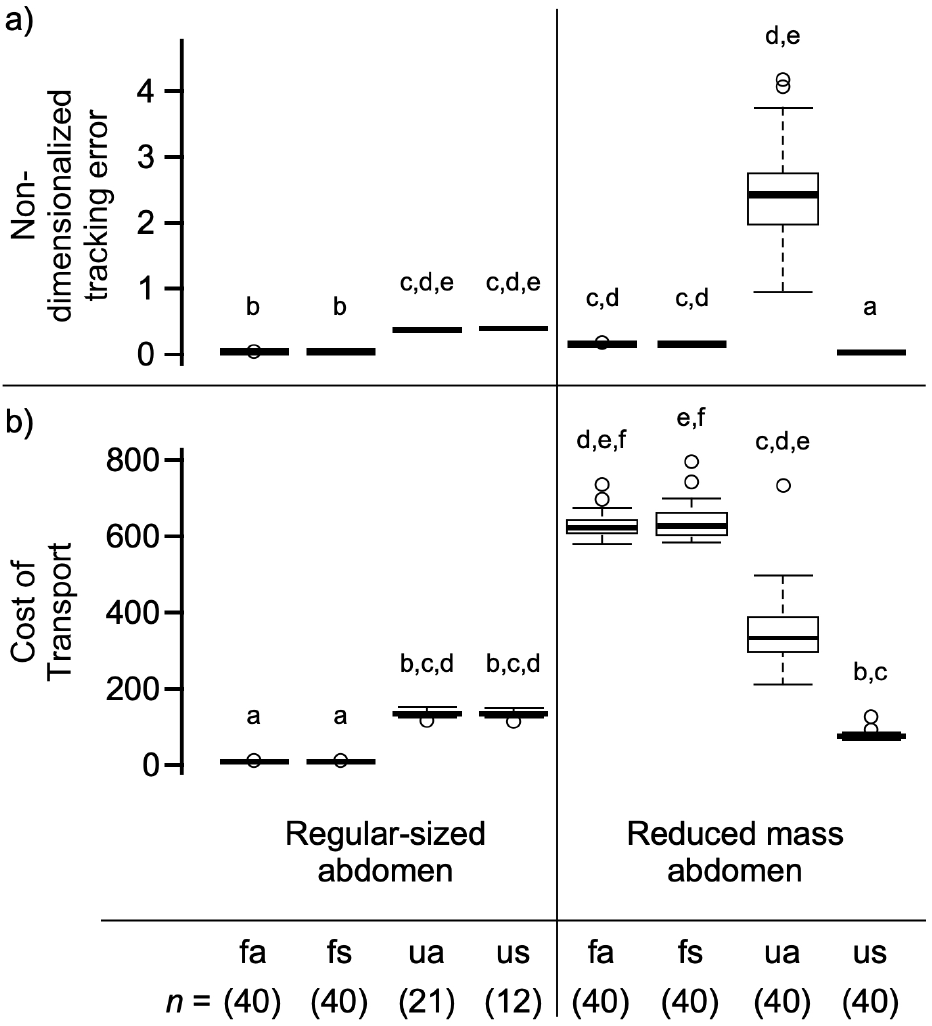
Panel (a) is the non-dimensionalized tracking error. Panel (b) is the cost of transport. The regular-sized abdomen is based off of our measured parameters (Appendix, Table S2), and the reduced mass abdomen is an abdomen mass decreased by ∼90% to evaluate the role of effectively eliminating the abdomen mass from the system. The grouping of four treatments on the left side of the panel correspond to the regular-sized abdomen, while the grouping of four treatments of the right side of the panel correspond to the reduced mass abdomen. For both panels, and both sizes of abdomen all treatment shorthands are defined as follows: fully actuated (“fa”), fully actuated and location of applied force shifted treatment (“fs”), underactuated (“ua”), and underactuated with location of applied force shifted treatment (“us”). All results here are based on 40 full simulations per treatment with the exception of the regular-sized abdomen underactuated simulations (“ua” and “us”). Each full simulation run time of 10 seconds of simulated flight. The letters above each box plot indicate statistical groupings. All statistical tests for significance were performed by a Kruskal-Wallis rank sum test. All statistical groupings were based on Bonferroni correction factor (*i*.*e*., reject hypothesis if *P* > 0.05/2). For non-dimensionalized tracking error, all model modifications except both fully actuated regular-sized abdomen treatments (left columns, “fa” and “fs”), and the reduced mass “us” treatment had significantly higher non-dimensionalized tracking error (*P* > 0.025). For cost of transport, all model modifications except both fully actuated regular-sized abdomen treatments (left columns, “fa” and “fs”) had significantly higher cost of transport (*P* > 0.025).

### Simulated flight: cost of transport

The cost of transport is defined as the non-dimensionalized work normalized to the weight of the head-thorax mass of the simulated insect multiplied by the distance it traveled in the 20 ms time frame. Cost of transport was lowest for the regular-sized abdomen fully-actuated simulations (Figure 3b, treatments “fa” and “fs”) and were significantly different from the other statistical groupings (Kruskal-Wallis *χ*^2^ = 295.27, *df* = 7, *P* = 2.2e-16, effect size, *η*^2^ = 0.935).

Energy expenditure was minimized for fully actuated treatments. Decreasing the abdomen mass by ∼90% yields flight dynamics similar to a normal abdomen (see Appendix, Figures S1, S2) for the underactuated and location of applied force shifted treatments. However, the cost of transport is higher for all reduced abdomen size treatments than the fully actuated, regular-sized abdomen treatments.

### Moth free flight: incidence of flight

One metric to observe any differences between the two treatments is the incidence of flight. After dark adaptation, moths were brought into the darkened arena and allowed 10 minutes to initiate flight. Flight or no flight was scored and compiled. Of 40 experimental treatment trials, 7 trials resulted in moth flight. Of 37 sham treatment trials, 18 resulted in moth flight. The incidence of flight is much lower in experimental treatment moths (17.5%) than in sham treatment moths (48.6%). A chi-squared test revealed that the incidence of flight in the experimental treatment is significantly lower than in the sham treatment (*χ*^2^ (1, *N* = 77) = 8.5053, *p* = 0.003541).

### Moth free flight: kinematics and path metrics

For all moths which successfully flew, we used various metrics to evaluate the dynamics of the flight paths, and observe the differences between the two treatments. These metrics included root mean square (RMS) velocity, RMS acceleration, tortuosity, box dimension, the log-log transform of the spatial frequencies of flight path, and flight duration.

The RMS velocity was not significantly different for both the experimental and sham treatments (Wilcoxon rank sum exact test, *P* = 0.6947). The RMS acceleration was not significantly different for both the experimental and sham treatments (Wilcoxon rank sum exact test, *P* = 0.6505). The sliding window tortuosity was not significantly different for both the experimental and sham treatments (Wilcoxon rank sum exact test, *P* = 0.261). The box dimension was not significantly different for both the experimental and sham treatments (Wilcoxon rank sum exact test, *P* = 0.6495). Neither the slopes nor intercepts of the log-log transform of the spatial frequencies of tortuosity were significantly different between experimental and sham treatments (Wilcoxon rank sum exact test, slope: *P* = 0.9408; intercept *P* = 0.3312).

However, the flight duration was significantly different for both the experimental and sham treatments (Wilcoxon rank sum exact test, *P* = 0.01561). This implies inhibition of abdominal movements yields shorter flight bouts for experimental treatment moths.

### Double-blind scoring of rolling maneuvers

The incidence of rolls in the experimental treatment was 45 of 96 (46.9%). The incidence of rolls in the sham treatment was 23 of 112 (20.5%). This difference was statistically significant (Wilcoxan rank sum exact test, *W* = 132, *P* = 0.01332). This implies inhibition of abdominal movements yields a greater incidence of rolls for experimental treatment moths, although it remains unclear if this is a behavioral compensation or an inertial consequence of the abdominal restriction.

## Discussion

In this study we have provided both theoretical and experimental evidence that abdominal motions can play a critical role in flight control. This is shown in both the inertial dynamics model of this study and in experimental treatments of freely flying hawkmoths. Three key results arise from our computational analysis. First, our computational approach, inspired by Model-Predictive Control (MPC), successfully predicts forces and torques required to accomplish the complex motor task of tracking a moving target. Second, we find that both fully actuated and underactuated models of control can successfully accomplish these tasks. However, the error associated with tracking is generally greater for underactuated systems. Third, we find that the energetic cost of transport, as measured by the energy divided by the product of the thorax muscle weight and distance traveled, depends quite strongly on the abdominal motions. We showed that all computational model variations were able to successfully track the vertically oscillating goal, yet two treatments in particular minimized the cost of transport (both fully actuated, regular-sized abdomen models, see Figure 3b, left columns, treatments “fa” and “fs”). We also showed that when abdomen motion was inhibited in freely flying moths attempting to feed from a stationary 3D printed flower, yielded a drastically lower incidence of flight, and for moths which successfully flew, the duration of the flight bouts were significantly shorter and less stable.

### Inverse solutions using MPC-inspired models

Flight control is a non-linear problem in which the inverse solution–finding forces and torques required to follow a specified path–can be quite challenging. As mentioned previously, a number of outstanding studies have solved this inverse problem using a variety of approaches. For example, Hedrick and Daniel [2006] used a genetic algorithm melded with a simplex optimization in which 10 parameters were used to solve the inverse problem of hovering and level flight control with the same four degrees of motion we allowed in our simulations. With 10 control parameters, they simulated an overactuated system, particularly given that the state space is dominantly four degrees of freedom. Nonetheless, that study showed itwas possible to fix all but two kinematic parameters and still accomplish successful flight (Hedrick and Daniel [2006], Table 2). Interestingly, that study also showed that a single underactuated kinematic parameter set could perform the task of hovering, albeit with considerably higher error. This particular underactuated kinematic parameter set was the inspiration for our study as underactuated systems have been of particular interest in synthetically designed engineered systems (Hinson et al. [2013], Mian and Daobo [2008]).

Additionally, Dyhr et al. [2013], created a model which was linearized about equilibrium points. They found that the model operates at the very edge of stability. In contrast, our model was intentionally not linearized and explored the number of actuators necessary for successful flight. That said, our current study did a simple linearization and finds very similar frequency dynamics emerge. Unlike the work by Dyhr et al. [2013], we did not explore specific frequency response dynamics to a range of forcing frequencies. Rather we sought to identify the underlying eigenfrequencies of the system.

Genetic algorithms have been used elsewhere in studies of flight control including but not limited to: identifying actuator placement, controller design, vehicle design to launch small satellites, and other general optimization processes (Steinberg and Page [1999]). While, to our knowledge, only Hedrick and Daniel [2006] incorporate the use of a genetic algorithm for computing the inverse problem of hovering in a biological context, a similar approach was used to compute wing kinematics in flight that minimized the energy required for hovering (Berman and Wang [2007]).

Analogously, MPC approaches for flight control have gained considerable attention in recent literature including on-board control of quadrotors (Bouffard et al. [2012]), target tracking in fixed wing aircraft (Stastny et al. [2017], Zanker [1988b]), and other synthetic systems (Shamaghdari et al. [2015]). However, to our knowledge, it has not been widely deployed as a method for exploring natural flight systems. MPC approaches also lend themselves to future studies in which deep neural networks (DNNs) can be developed, refined, and influence the basis of problems with otherwise infinite state-space permutations. In summary, the implementation of MPC approaches coupled with DNNs can provide unique insight into a vast array of physically bounded, biologically relevant questions.

### Fully actuated and underactuated controls

For our model of a flying moth moving with four degrees of freedom–in our cases two translational (*x, y*) and two rotational (*θ, ϕ*) directions–we explored two control scenarios, one in which the model is fully actuated using four control inputs, and one in which it is underactuated, where we use just three control inputs. Additionally, we examined the role implicit torques (“fs” and “us”) may play with regard the flight performance metrics described in this study. Formally, underactuated systems are those for which the control system cannot accomplish any acceleration in some of the degrees of freedom at any instant in time (Shkolnik and Tedrake [2008]). Underactuated systems have received considerable attention in recent years, often because they are commonly seen in a variety of biological systems. For example, Xinyan Deng et al. [2006] show that time invariant averaging theory can be used for controlling bioinspired insect robots that are underactuated. Similar approaches have been used for underactuated fish-like robots (Morgansen et al. [2001]).

Similar to the time-averaging methods above, in our underactuated scenario, we use a fixed force and fixed abdominal torque during the equivalent time of half a wing stroke to effect control, effectively serving as a stroke averaged control. As we noted above, this approach leads to effective tracking, though an intriguing trade-off appears. When the abdominal mass is close to that observed in *Manduca* the cost of transport in both underactuated configurations (“ua” and “us”) is greater than that associated with the fully actuated system (“fa” and “fs”) (Figure 3b). As observed, if the abdominal mass is drastically smaller, as in our reduced mass abdomen models, the cost of transport is actually greater for the fully actuated system despite the dynamics yielding reasonable motions (Figure S2). The cost of transport of underactuated systems for both the normal abdominal mass and the reduced mass are similar in magnitude (Figure 3b). If the role of the abdomen is nearly completely removed (*i*.*e*. mass reduced by ∼90%), this suggests that an underactuated system may be best to perform the task of tracking a vertically oscillating signal. This is because the reduced mass abdomen does not require more actuators to stabilize the motion. This result may inform the development of micro air vehicles carrying significantly smaller loads (*i*.*e*. quad-copters). Furthermore, the model developed in this study allows for the exploration of different abdominal sizes and spatial configurations in future simulations.

When considering the role of implicit torques, we observed that the fully actuated simulations (regardless of abdomen size) yield statistically similar flight performance metrics (Figure 3). This trend remains consistent for the underactuated simulations only when the abdomen was regular-sized. Yet when the role of the abdomen was nearly completely eliminated (*i*.*e*. mass reduced by ∼90%), the underactuated scenarios benefited from having a shifted location of applied force (“us”) despite using one fewer control input. This implies such underactuated models with reduced abdominal mass would benefit from an implicit torque on the system.

Taken together, the flight dynamics (Appendix, Figures S1, S2), non-dimensionalized tracking error (Figure 3a), and cost of transport (Figure 3b) yield conflicting results with the findings of Hedrick and eDaniel [2006] (see Hedrick and Daniel, Table 2). For the normal mass simulations, a higher non-dimensionalized tracking error, and cost of transport was shown for both underactuated treatments (“ua” and “us”) when compared to the fully actuated treatments (“fa” and “fs”) which is in accordance with Hedrick and Daniel [2006]. However, the flight dynamics (Appendix, Figures S1, S2) reveals that while the model can track rectilinear motions accurately (*x, y*), and even maintain a relatively reasonable head-thorax rotation (*θ*, with some abrupt jostling), the abdominal motions spin indefinitely for both underactuated configurations (“ua” and “us”). This dramatic abdominal spin demonstrates a conflict with the findings of Hedrick and Daniel [2006].

In all instances when applied wing torques or offset origin of wing forces are absent from the system, we observe unstable dynamics (“ua” treatment in Appendix, Figures S1, S2), high energy expenditure, and high tracking error (Figure 3, “ua” models). This implies the necessity of an applied torque, regardless of how miniscule, is essential for dynamical stability. Interestingly, the pitch moment of inertia about the center of mass for the entire head/thorax and abdomen system will depend on abdominal flexion angle: greater flexion has a lower pitch moment of inertia.

### Restricting natural abdominal motions decreases incidence of flight and flight duration

For freely flying moths given the task of feeding from a stationary 3D printed flower, our sham treatment moths flew successfully with no discernible inhibition to their flight pattern. Flight incidence was dramatically decreased for moths given this same task with the carbon fiber rod restricting their abdominal motion. Additionally, if moths flew, various flight dynamics (*e*.*g*. velocities, path tortuosity) were not significantly different between treatments, except for length of flight duration. This indicates the experimental treatment moths that were able to fly were inhibited to the point of having shorter flight bouts. As suggested by Figure 3, the shorter flight bouts may indeed be a result of the increased cost of transport because of the reduced number of actuators.

While simple metrics of flight dynamics such as speeds, accelerations, or tortuosities did not yield significant differences between moths with restricted abdominal motions and control individuals, moths in the experimental treatment group were observed to engage in a higher incidence of rolling motions than sham treatment moths. We suspect that abdominal flexion could increase the rotational moment of inertia in the roll axis: a straight cylinder has a lower roll moment of inertia than a bent cylinder of identical length and mass. Indeed, maneuvering via abdominal flexion may allow roll motions when straight and roll stability when bent.

Similarly, this inhibition of abdominal movements is shown to have adverse effects on other animals as seen in a number of other studies. For example, juvenile mantids that had their abdominal segments restricted by superglue resulted in slower rotations necessary to complete a jump (Burrows et al. [2015]), sometimes causing them to crash head-first into their task goal–an otherwise easy to accomplish task for normal juvenile mantids. Live wingless aphids assume a stereotypical body posture before falling off leaves, and an inertial mechanism aerially rights their positions during free fall (Ribak et al. [2013]). However, this righting mechanism was significantly reduced when the wingless aphids were dead or had ablated appendages. At critical stages of aerial righting, wingless nymphal stick insects actively use their legs to reorient while falling (Zeng et al. [2016]). Gliding ants (wingless ant workers) apply active control over their aerodynamic forces when righting their path back to tree trunks (Munk et al. [2015]) despite their lack of having aerodynamically streamlined appendages to accomplish this task.

Future work could include 3D motion tracking of moths using the same vertically oscillating signal in our MPC-inspired model on live moths. This potential new study could also include varying degrees of inhibited abdominal motion ranges (*i*.*e*., potential halfway degrees of treatment–sticks which get stiffer as flexion increases).

## Conclusion

Airframe deformation: inertial reconfiguration contributes to flight control Taken together, both the computational and experimental results point to an important role for inertial reconfiguration of body elements in movement control. While wings are clearly the most important structures for animal flight control, this study shows that inertial components of non-wing structures also play an important role.

## Appendix

### Model formulation and methods

The Euler-Lagrange equations form a system of four coupled second order ordinary differential equations for the state variables *q*, (*q* = *x, y, θ, ϕ*).

The Euler-Lagrange equation is written in the following general form:

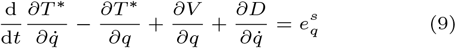

Where *q* is a placeholder for a flow variable (*x, y, θ, ϕ*). *x* and *y* are with respect to the standard Cartesian coordinates, *θ* is the angle of the head-thorax mass with respect to the x-axis, and *ϕ* is the angle between the midline of the abdomen with respect to the x-axis (Figure 1b).

Furthermore, *T* ^∗^(*p*) is defined as the kinetic co-energy, V is defined as the potential energy, *D*(*p*) is defined as the dissipative energy, and 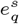 the external effort exerted that affects the respective flow variable (either *x, y, θ, ϕ*). This yields four Euler-Lagrange equations for the four variables of this model:

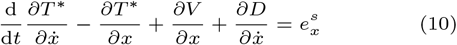

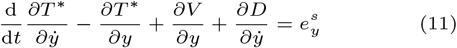

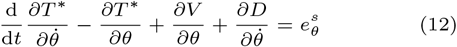

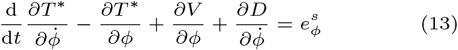

The positional vectors of our two masses with respect to the thorax-abdomen joint are the following:

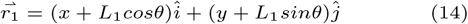

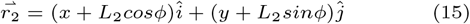

Where *L*_1_ is a fixed length between the thorax-abdomen joint to the center of the head-thorax mass (Figure 1b). Similarly, *L*_2_ is a fixed length between the thorax-abdomen joint to the center of the abdomen mass (Figure 1b).

The velocity vectors for each of the two masses are simply the derivatives of the positional vectors with respect to time as follows:

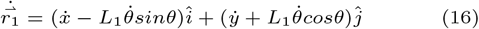

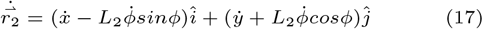

The kinetic energy of the system is as follows:

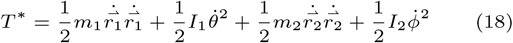

Where *m*_1_ is the mass of the head-thorax, *I*_1_ is the moment of inertia of the head-thorax mass, *m*_2_ is the mass of the abdomen, and *I*_2_ is the moment of inertia of the abdomen. When the velocity vectors are substituted in, (18) becomes the following form:

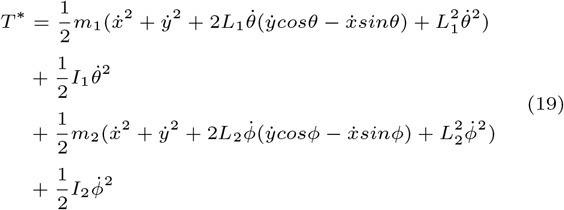

The potential energy of the system is as follows:

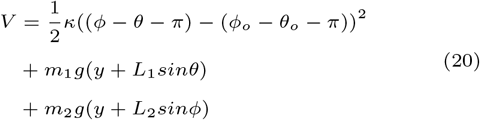

Where *κ* is the torsional spring constant of the joint that connects the thorax to the abdomen, *g* is the acceleration due to gravity, *ϕ*_*o*_ is the initial abdomen angle, and *θ*_*o*_ is the initial head-thorax angle. And *β*_*rest*_ is defined as: *ϕ*_*o*_ -*θ*_o_ *π*.

The dissipative energy of the system is as follows:

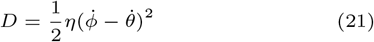

Where *η* is the torsional damping constant of the joint that connects the thorax to the abdomen. The work done by the applied efforts to the system are as follows:

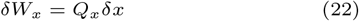

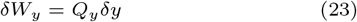

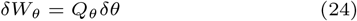

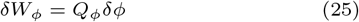

Therefore, the applied efforts are as follows:

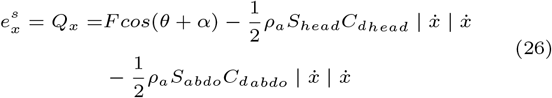

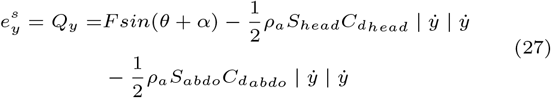

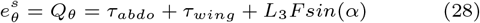

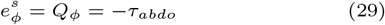

Where *F* is the averaged aerodynamic lift force over the course of the 20 ms simulation time period caused by the flapping wings of the insect, *α* is the angle of the aerodynamic lift force with respect to the midline of the head-thorax mass, *ρ*_*a*_ is the density of air, *S* is the surface area of the insect which is experiencing aerodynamic drag (modeled as a sphere).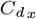, and 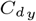 are the coefficients of drag in the x-direction and in the y-direction respectively, *τ*_*abdo*_ is the torque applied to the thorax-abdomen joint for flight correctional purposes and/or to counteract external perturbations (note: conservation of angular momentum requires an equal and opposite torque applied by the abdomen), and *L*_3_ is the fixed length between the thorax-abdomen joint to the center of the aerodynamic lift of the head-thorax mass.

Taking the appropriate derivatives of equations 19-21 as required by equations 10-13, yield the following four equations of motion (note, not all derivatives mentioned in equations 10-13 are necessary):

Equation of motion in the x-direction:

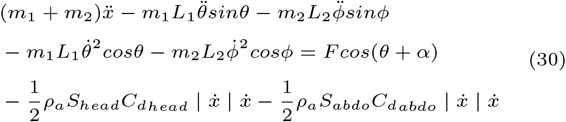

Equation of motion in the y-direction:

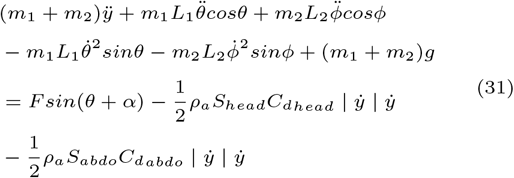

Equation of motion in the theta-direction:

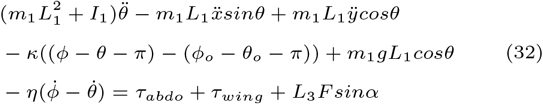

Equation of motion in the phi-direction:

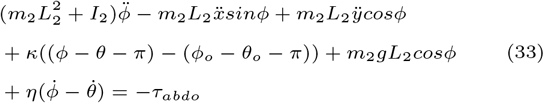

These equations of motion (Equations 30 - 33) are the ones used in our inertial dynamics model.

### Fixed mechanical properties

The mechanical properties of the model are fixed throughout the simulations and are defined in Table S2. The terms for mass and moment of inertia in Figure 1 are defined as follows using the terms in Table S2.

Mass:

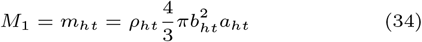

Moment of inertia:

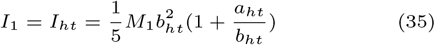

Where the subscript “*ht*” is shorthand for “head-thorax.” Similarly, the mass and moment of inertia of the abdomen correspond to *M*_2_ and *I*_2_ respectively.

### Loss function weighting coefficients

The loss function weighting coefficients are defined in Table S3. These coefficients ensure that deviations from our goal position and rotation heavily penalize rectilinear motion deviations (*x, y*), with head motion deviations penalized secondarily as strong (*θ*). The derivatives of these motions 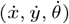 are not as strongly penalized.

**Table S1.** The prime number frequencies (in Hz) listed out in sequential order and their respective amplitudes (in cm). Amplitudes decreased with increasing frequency to prevent the system from going unstable.

| Goal motion<br>Frequency (Hz) | Goal motion<br>amplitude (cm) |
| --- | --- |
| 0.2 | 5 |
| 0.3 | 3.3333 |
| 0.5 | 2 |
| 0.7 | 1.4286 |
| 1.1 | 0.9091 |
| 1.7 | 0.5882 |
| 2.9 | 0.3448 |
| 4.3 | 0.2326 |
| 7.9 | 0.1266 |
| 13.7 | 0.0730 |
| 19.9 | 0.0503 |

### Prescribed goal motion frequencies and amplitudes

The frequencies of the prescribed goal motion are 11 prime number frequencies drawn directly from Roth et al. [2016]. The purpose of these prime number frequencies is two-fold: 1) to determine if the output of the system is linear or non-linear, 2) to ensure that any potential harmonics of the system can be distinguished from their basal frequencies (*i*.*e*. not overlap).

The amplitude(s) of the goal motion signal decrease by a prescribed factor as shown in equation 36. This amplitude decrease is necessary to ensure that the derivative of the models tracking this goal do not accelerate substantially causing unwanted instabilites in the system.

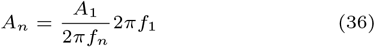

**Table S2.**
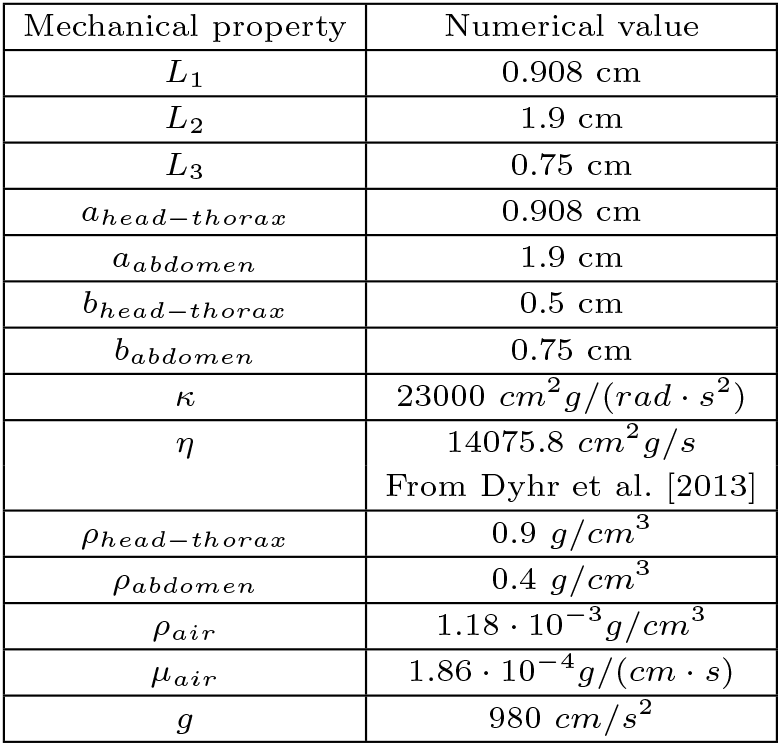
The mechanical properties of the model are fixed throughout the duration of all simulations. All values were calculated from our own measurements and rounded as appropriate with the exception of one: the torsional damping coefficient (*η*) which was previously measured in Dyhr et al. [2013].

| Mechanical property | Numerical value |
| --- | --- |
| $L_1$ | 0.908 cm |
| $L_2$ | 1.9 cm |
| $L_3$ | 0.75 cm |
| $a_{head-thorax}$ | 0.908 cm |
| $a_{abdomen}$ | 1.9 cm |
| $b_{head-thorax}$ | 0.5 cm |
| $b_{abdomen}$ | 0.75 cm |
| $\kappa$ | $23000 \text{ cm}^2 \text{g}/(\text{rad} \cdot \text{s}^2)$ |
| $\eta$ | $14075.8 \text{ cm}^2 \text{g}/\text{s}$<br>From Dyhr et al. [2013] |
| $\rho_{head-thorax}$ | $0.9 \text{ g}/\text{cm}^3$ |
| $\rho_{abdomen}$ | $0.4 \text{ g}/\text{cm}^3$ |
| $\rho_{air}$ | $1.18 \cdot 10^{-3} \text{ g}/\text{cm}^3$ |
| $\mu_{air}$ | $1.86 \cdot 10^{-4} \text{ g}/(\text{cm} \cdot \text{s})$ |
| $g$ | $980 \text{ cm}/\text{s}^2$ |

**Table S3.**
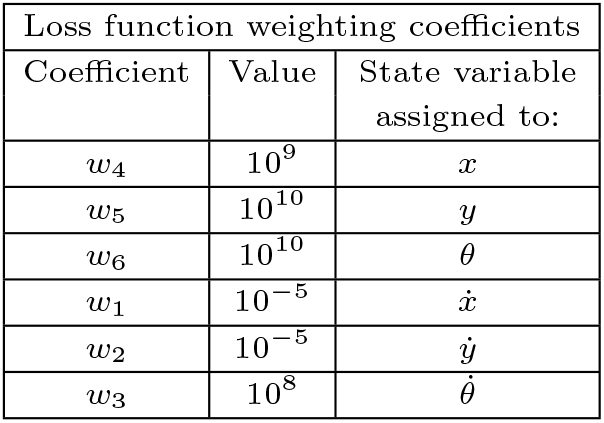
The weighting coefficients of the loss function penalize rectilinear motions higher than head-thorax motion. Such weights are critical for the loss function to function.

| Loss function weighting coefficients |  |  |
| --- | --- | --- |
| Coefficient | Value | State variable<br>assigned to: |
| $w_4$ | $10^9$ | $x$ |
| $w_5$ | $10^{10}$ | $y$ |
| $w_6$ | $10^{10}$ | $\theta$ |
| $w_1$ | $10^{-5}$ | $\dot{x}$ |
| $w_2$ | $10^{-5}$ | $\dot{y}$ |
| $w_3$ | $10^8$ | $\dot{\theta}$ |

### Derivation of torsional spring constant system

As stated in Methods, the moths used in this experiment can be modeled as a system of two masses (thorax and abdomen) connected with a torsional spring constant and a torsional damper (Figure 1b), and represented by Equation 37

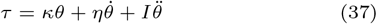

Which can be transformed into Equation 38

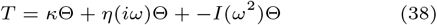

This can be rearranged into a torque-angle ratio form as seen in equation 39

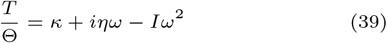

The system yields two components: the real component (Equation 40), and the imaginary component (Equation 41).

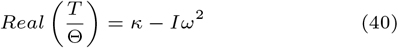

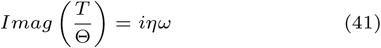

The real component corresponds to the torsional spring constant and can be rearranged into Equation 8 (see Methods)to calculate the torsional spring constant. The imaginary component corresponds to the torsional damping coefficient which is not calculated in this study, but rather, a value previously calculated by Dyhr et al. [2013] is used.

### Measurements of body segment masses

Five male hawkmoths, and five female hawkmoths had their total mass, head, thorax, and abdomen weighed. For a full distribution of these values, see Table S4.

### Estimating the fundamental frequency

The set of nonlinear second order differential equations (Equations 30-33) represent a dynamical system that has effective masses, damping components and stiffness values. To explore the fundamental frequencies that underlie this system we linearized our system of equations for a moth that is hovering. We set the position of the moth to be the origin and the angle of the body to be *pi/*4 with the abdomen aligned with the head-thorax:

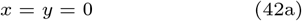

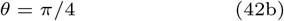

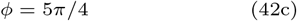

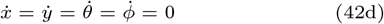

We linearized the system about these conditions and solved for the mass matrix *M*, the damping matrix *C* and the stiffness matrix *K* operating on the state vector *q* = [*x, y, θ, ϕ*] and its derivatives such that:

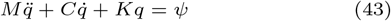

where *ψ* is the vector of applied forces and torques. In the limiting case of zero damping we solve the eigenvalue problem:

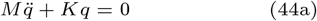

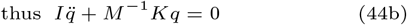

Setting 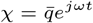 we find arrive at the eigenvalue problem:

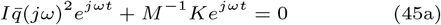

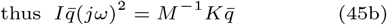

where *I* is the identity matrix and the fundamental frequencies are (*jω*)^2^. Given the parameters laid out in Table S2, we find the lowest fundamental frequency to be 1.99 Hz for the case of hovering. This linearized estimate of the modal frequency is quite close to the frequencies we observe in Figure S1. As such, the oscillations that result follow from the dynamics of this coupled system. Details of the parameter values and the structure of the matrices *M, C*, and *K* can be found in our Github repository in a file Modes.nb.

## Competing interests

No competing interests.

**Table S4.** Body segment mass distribution in grams. The mean and standard deviation values are calculated of the set of five male moths and five female moths respectively. The minimum and maximum of each set of five male moths and five female moths are also reported. The parenthetical values report the corresponding percentages of the value within the cell. Note the percentages summed across rows do not sum to 100% because 1) the wings and legs were not included in this measurement, and 2) the distribution reported reflects the value of the set (*i*.*e*., the minimum/maximum values not be shared by the same individual moth).

| Sex -<br>Statistic | Total<br>mass<br>(g) | Head<br>mass<br>(g) | Thorax<br>mass<br>(g) | Abdomen<br>mass<br>(g) |
| --- | --- | --- | --- | --- |
| *Male*<br>*average* | 2.0758 | 0.135<br>(6.5%) | 0.584<br>(28.3%) | 1.0252<br>(49.3%) |
| *st.dev* | 0.145 | 0.0267 | 0.0449 | 0.126 |
| *min.* | 1.953 | 0.100<br>(5.1%) | 0.532<br>(24.9%) | 0.916<br>(46.0%) |
| *max.* | 2.302 | 0.173<br>(7.5%) | 0.651<br>(32.7%) | 1.187<br>(53.1%) |
| *Female*<br>*average* | 3.550 | 0.183<br>(5.2%) | 0.686<br>(19.6%) | 2.206<br>(61.9%) |
| *st.dev* | 0.489 | 0.0264 | 0.0692 | 0.394 |
| *min.* | 2.790 | 0.148<br>(4.2%) | 0.618<br>(15.8%) | 1.660<br>(59.3%) |
| *max.* | 4.054 | 0.214<br>(6.2%) | 0.796<br>(22.7%) | 2.728<br>(67.3%) |

## Author contributions statement

Intentionally left blank.

## Acknowledgments

Intentionally left blank.

---

**Fig. S1.**
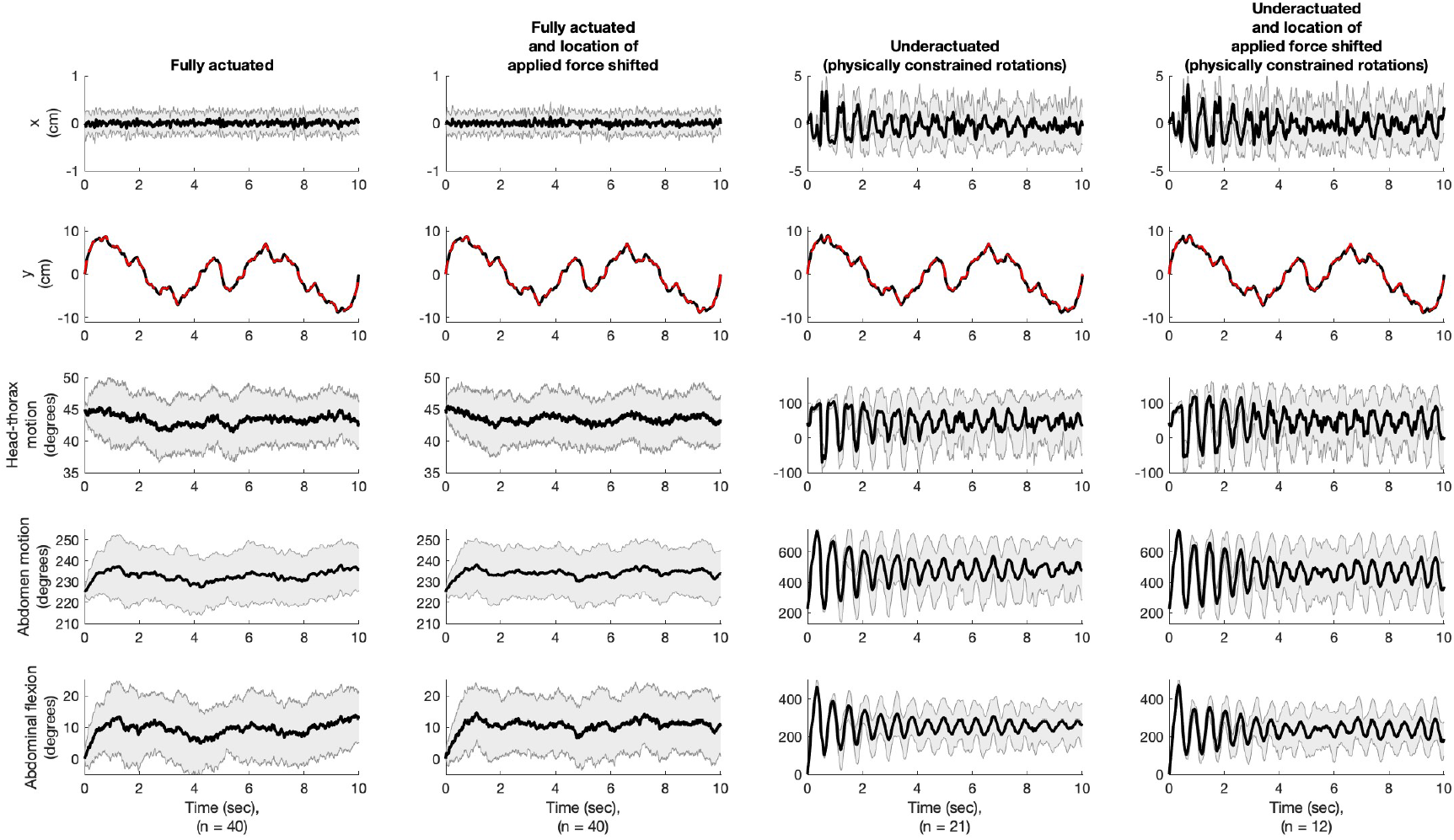
Model dynamics output for all model variations of the regular-sized abdomen model. All plots are with respect to time for a 10 second simulation period. Top row is the x-motion in cm. Second row is the y motion in cm (note the dashed red line is the goal motion of our vertically oscillating goal). The third row is the head-thorax (*m*_1_) mass motion in degrees. The fourth row is the abdominal motion (*m*_2_) in degrees. The bottom row is the abdominal flexion angle (*i*.*e*. difference between rows 4 and 3 respectively) in degrees. For all treatments, the model tracks the input y-motion well.

**Fig. S2.**
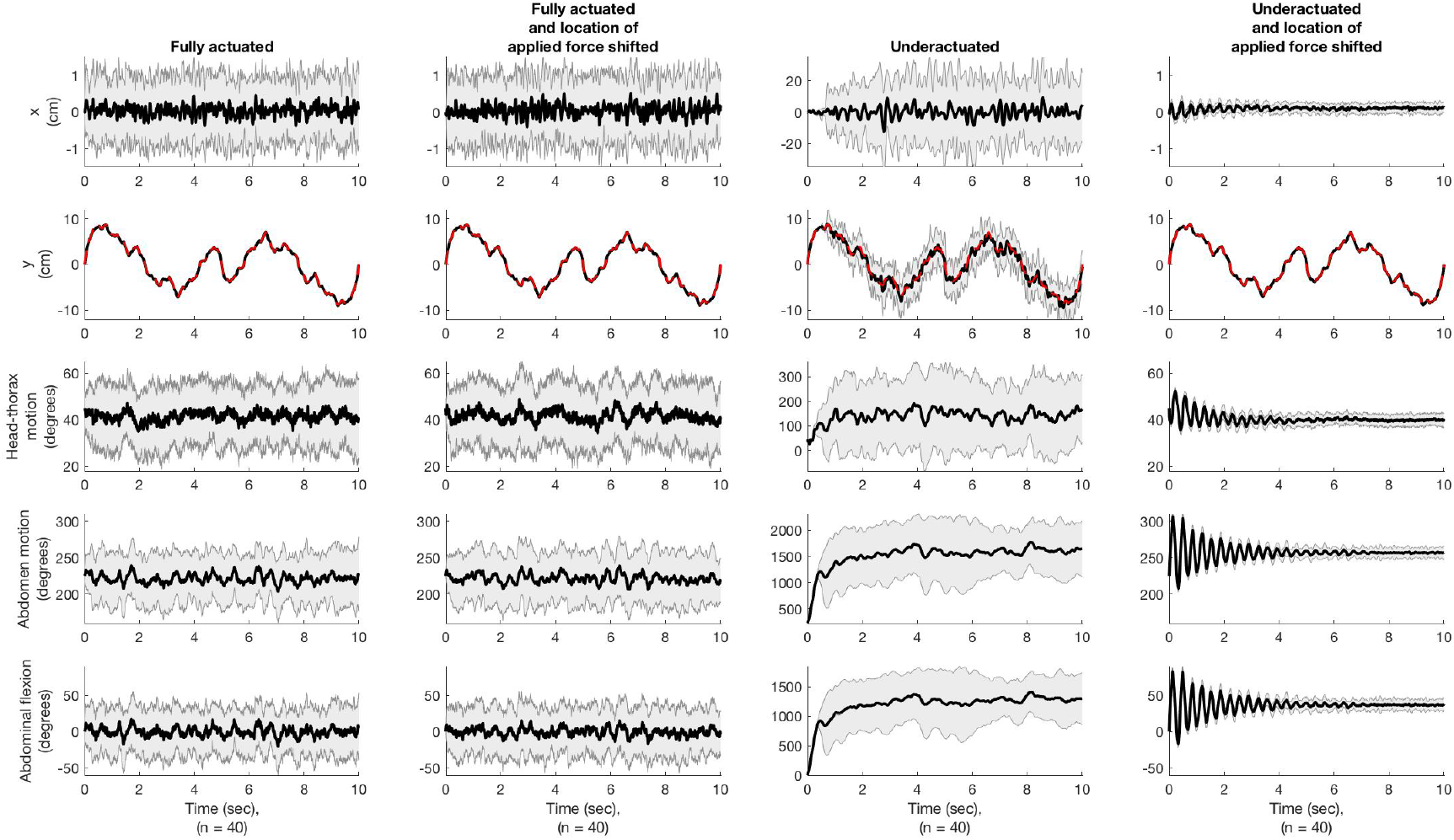
Model dynamics output for all model variations of the reduced-sized abdomen model. All plots are with respect to time for a 10 second simulation period. Top row is the x-motion in cm. Second row is the y motion in cm (note the dashed red line is the goal motion of our vertically oscillating goal). The third row is the head-thorax (*m*_1_) mass motion in degrees. The fourth row is the abdominal motion (*m*_2_) in degrees. The bottom row is the abdominal flexion angle (*i*.*e*. difference between rows 4 and 3 respectively) in degrees. For all treatments, the model tracks the input y-motion well.

## Author Name

This is sample author biography text. The values provided in the optional argument is meant for sample purpose. There is no need to include the width and height of an image in the optional argument for live articles. This is sample author biography text this is sample author biography text this is sample author biography text this is sample author biography text this is sample author biography text this is sample author biography text this is sample author biography text this is sample author biography text.

## Author Name

This is sample author biography text this is sample author biography text this is sample author biography text this is sample author biography text this is sample author biography text this is sample author biography text this is sample author biography text this is sample author biography text.

